# A depositional taphonomy model for sedaDNA based on biotic profiles of contrasting sedimentation regimes

**DOI:** 10.1101/2025.04.30.651409

**Authors:** Teri A Hansford, Vivian Higgs, Vincent Gaffney, Logan Kistler, Robin G Allaby

## Abstract

Sedimentary ancient DNA (sedaDNA) has become an important tool in Quaternary Science, but still little is understood of its taphonomy and whether sedaDNA represents the local vicinity or originates from distant sources. Here we show key insights can be made about the origins of sedaDNA by integrating sedimentological and sedaDNA data into a sediment influx depositional model. We reconstruct contrasting taphonomic regimes of lacustrine, alluvial, terrestrial and permafrost systems. Lacustrine systems show signals of regional catchment sources of sedaDNA, while alluvial, terrestrial and permafrost systems show a greater influence of DNA from the immediate local environment. However, there is a prevalence of mixed ecosystem signals which may span a broad temporal timeframe if sediment reworking has occurred. The ability to distinguish between local and distal sources of sedaDNA is of major importance in the interpretation of taxonomic profiles to reconstruct palaeoenvironments.

## Introduction

Over the past two decades sedimentary ancient DNA (sedaDNA) has revolutionized the resolving power of Quaternary science^1,2^, from the identification of a forested Greenland prior to the last glaciation^3^ to more recently the recovery of unexpected ecosystem profiles over 2 million years ago^4^. Correlates between environmental factors and DNA damage suggest that DNA may persist on the million-year time scale to a degree largely determined by temperature and temperature fluctuation^5^, followed by other factors such as chemistry^6^ and physical adsorption to protective mineral surfaces^7^. However, concerns were raised early in the field about taphonomic processes which could influence DNA deposition leading to erroneous interpretations of sedaDNA^8^.

There are a number of taphonomic factors when interpreting sedaDNA profiles that should be considered. Among these, there may be biases in the amount of DNA laid down as a consequence of organism size^2^ and genome size^9^ leading to over or under representation of species. Secondly, post deposition movement of DNA through potential diffusion^10^, leaching^8^ or sediment reworking^11^ can lead to anachronous signals. Thirdly, considerations are required concerning the source of DNA deposited in a sediment^12^. Sediments are usually laid down under fluids originating from a source environment, which may be some distance from the deposition environment and may well not reflect the local environment around the site of deposition. It is possible that sedaDNA derives from such incoming sediment, effectively representing a case of reworking or a general catchment signal, or alternatively it may originate from the local environment being laid down concurrently with incoming sediment. In these scenarios, a sedaDNA signal could be reflective of a local, distal or regional signal, leading to different sets of interpretations. Several studies have reported an apparently local source of sedaDNA^13–16^, while there have also been recent attempts to build taphonomy models which incorporate the influence of catchment signal resulting from sedaDNA being brought with influxing sediments from surrounding regions^12^. There is therefore a need to integrate sedimentological data with sedaDNA data to better understand depositional sources of sedaDNA.

### A depositional framework integrating sediments and DNA

If the presence of sedaDNA is tightly linked to sediments, then the sedimentation regime should be highly influential on sedaDNA profiles. Here, we describe and test a framework of a depositional model that measures the influence of sediment profiles on sedaDNA profiles, facilitating a measure of the influence of local and influxing sources of DNA on sedaDNA profiles.

In a depositional environment, when the type of sediment being deposited changes, it likely reflects some change in the source environment from which the sediment originates. This could be a physical change in locality from which the sediment originates, or some other large change in that source environment which also could be associated with biotic turnover. If sedaDNA is directly associated with sediment also originating from the source environment, then sedaDNA profiles should be heavily influenced by sediments. However, if sedaDNA is predominantly from local sources independent of the sediment, then the sedimentation regime may be expected to be less influential on sedaDNA profiles. Recently, a framework integrating sedaDNA profiles and sedimentological data was developed on the basis of these assumptions and applied to marine core samples from the Doggerland area^11^, Figure 1. In this model the amount of biotic change between two time points is considered under two different sedimentation scenarios. The change in taxonomic profile, as evidenced in sedaDNA from one point in time to the next, is assumed to be a function of the amount of taxonomic change that has occurred in the local and the distal source environments respectively, and their relative contributions to the sedaDNA profile. By considering two scenarios, one in which there has been no change in sedimentation regime (distance value ∂1), and a second in which there is such a change (distance value ∂2), several parameters are fitted under two complex simultaneous circumstances. Parameter fitting is achieved through a Monte Carlo approach to find values of parameters producing ∂1 and ∂2 that most closely match the observed measurements (see Methods).

**Fig. 1.**
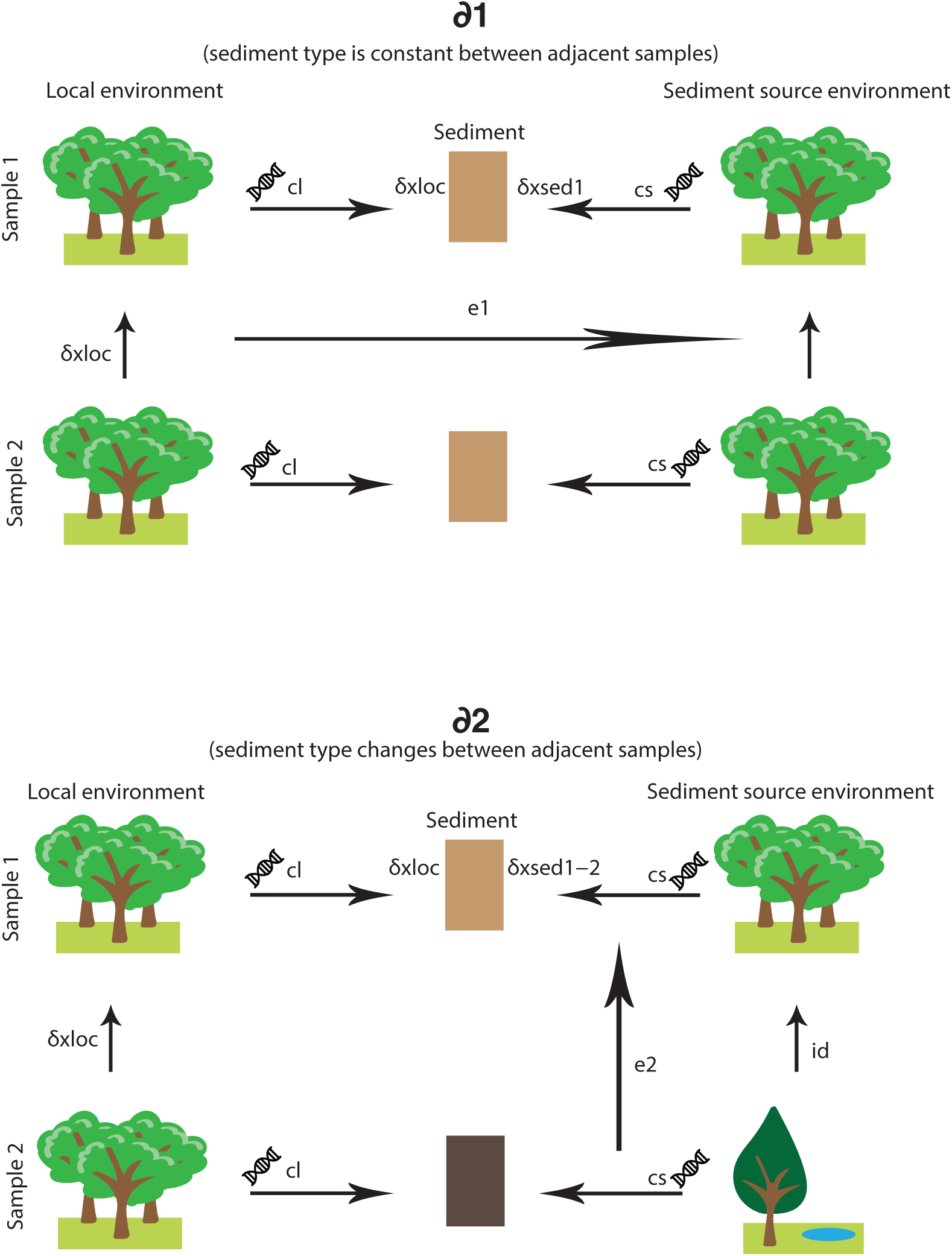
Sediment influx depositional model. TOP. ∂1 scenario: adjacent samples of same sediment type 1. BOTTOM. ∂2 scenario: adjacent samples of sediment types 1 and 2. Parameters as follows: ∂1 – ecological distance between adjacent samples of the same type; ∂2 – ecological distance between adjacent samples of different sediment types; cl – proportion of sedaDNA contribution from local sources; cs – proportion sedaDNA contribution from influxing sediment from source environment; δ_loc_ – change in ecological profile of local environment over time; δ_xloc_ – contribution of δ_loc_ to ecological distance ∂ between samples; δ_sed1_ – change in ecological profile of influxing sediment source environment over time; δ_xsed1_ contribution of δ_sed1_ to plant ecological distance ∂ between samples; id – ecological distance between different influxing sediment source environments 1 and 2; δ_xsed1-2_ – contribution of id to ecological distance ∂ between samples; e1 – environmental variable 1 describing the difference in rate of ecological profile change over time between 8loc and 8sed1; e2 – environmental variable 2 describing the difference in influx rates from source environments 1 and 2.

This approach demonstrated that in a riparian (alluvial) system fine silt or clay sediments, indicating slow flow rates, were associated with high influence of local contribution and low levels of influxing sedaDNA from distal sources^11^. Conversely, coarse sand or gravel sediments, indicating more rapid flow rates, were associated with a high influence of influxing sedaDNA and a tendency toward mixed ecosystem and sediment reworking signals. Intuitively, such results make sense given the short half-life of DNA in water^6^, indicating that sedaDNA does not persist in significant amounts relative to local DNA sources in slow moving systems.

However, this framework assumes that changes in sedimentation regime are not generally correlated with wider climatic changes and regional biome turnover. In such a case, changes in sediment regime would be associated with variations in taxonomic profile both locally and distally, thereby causing a spurious correlation between sedimentation regime and taxonomic change. The sediment influx depositional model also does not take account of the possibility that different sediments may influence sedaDNA profiles through variant adsorbent properties^7^, which could cause DNA associated with particular sediment types to become over-represented so invalidating estimates of contribution. Furthermore, the general applicability of the framework is unclear since it has not been applied to non-riparian systems, such as lacustrine and terrestrial systems. Here, we test the robustness of the underlying assumptions of the framework and assess its general applicability to sedaDNA studies.

## Results

### Environmental associations of sediment regime and sedaDNA damage

To test the underlying assumptions of the depositional model framework shown in Fig. 1., we utilized the data set generated from Doggerland^11^ as this represents the most comprehensive assembly of sediment types and associated sedaDNA profiles to date, and spans a timeframe from the Pleniglacial to the late Holocene (>16kya-4Kya) encompassing several major climatic shifts. This is also the data set on which the framework was originally applied allowing us here to further interrogate the original model application and inferences. Firstly, we considered whether rates of sedimentation changes correlated with known major climatic shifts, Fig. 2a. While there is fluctuation apparent in sediment turnover, it is not notably different during events including the Younger Dryas and Pleistocene/Holocene boundary to later in the Holocene. The biotic profile shows some increase in the rate of change beginning at the Pleistocene/Holocene boundary, but shows the largest change abruptly around the time of the 8.2K cooling event^17^, which in this data suggests a profound change in the Doggerland area and a general expected profile of taxonomic change associated with broad climatic trends. The rate of sediment change does not seem to coincide with climatic events, or indeed broadly taxonomic change. The underlying assumption of the depositional model that sediment change is not associated with general climatic change and associated biotic turnover seems to be satisfied in this data set. In this case changes in sedimentation regime appear to be dominated by ongoing local processes.

**Fig 2.**
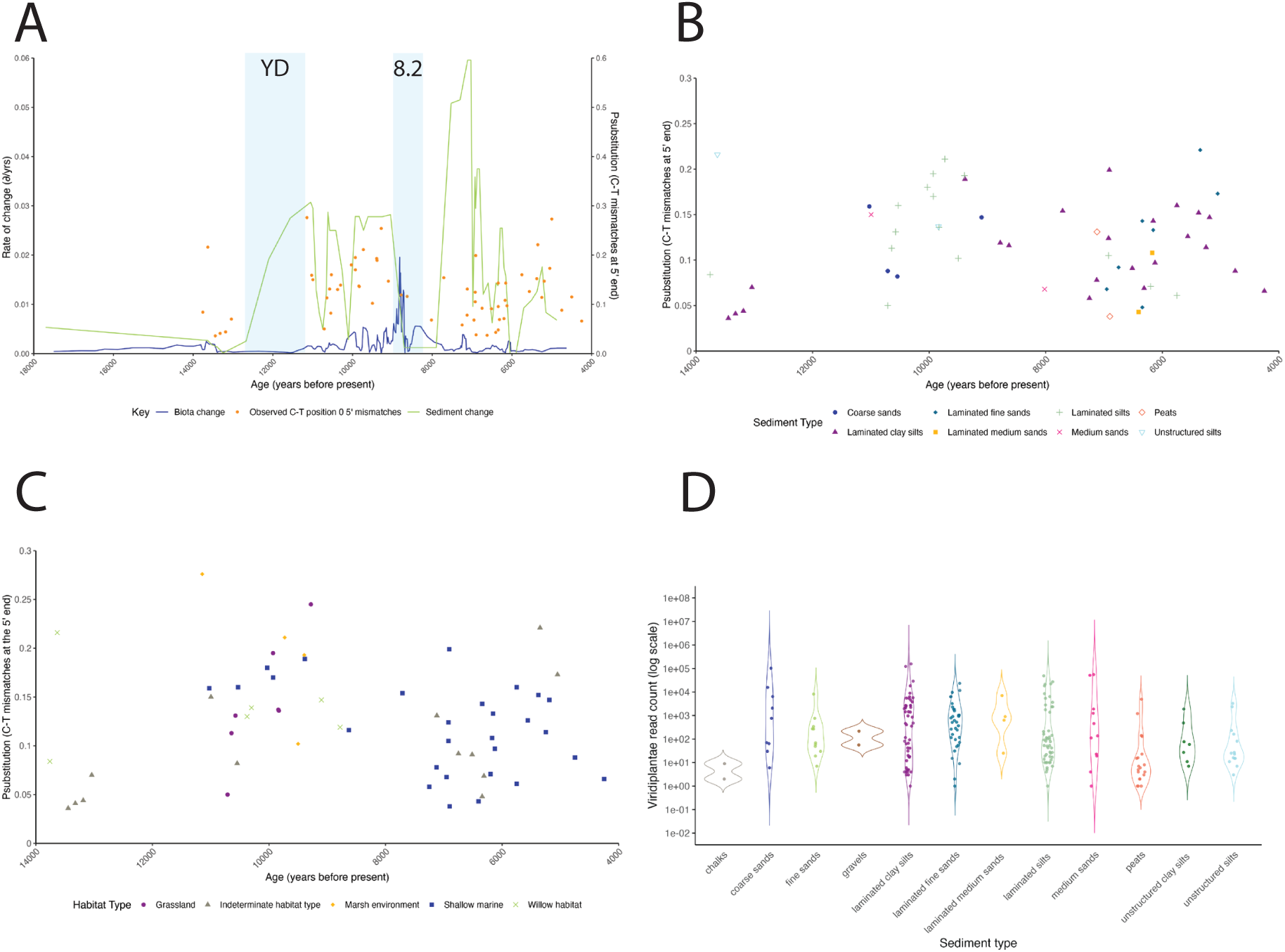
Environmental associations between sediment and sedaDNA. A. Rates of change in sedaDNA and sediments over time. Sliding window average of ecological difference (∂) per year (blue) and number of sediment changes per year (green) between adjacent samples. Observed C to T mismatch frequencies of individual samples shown in orange. Younger Dryas (YD) and the 8.2 ka event (8.2) are marked by blue boxes. B. Observed C to T mismatch frequency at position 0 of samples by sediment type (key under panel). C. Observed C to T mismatch frequency at position 0 of samples by dominant habitat type (key under panel). D. Filtered read number of Viridiplantae per sample by sediment type.

We assessed the influence of sediment chemistry on sedaDNA profiles. If DNA is adsorbed to surfaces of minerals in sediments, then its availability to hydrolytic attack should be reduced^7,18–21^. Expected consequences of this include a higher retention of DNA, greater persistence of DNA and a lower DNA damage profile associated with adsorbing sediments^22^. We examined the extent of DNA 5’ cytosine deamination at the terminal most position of DNA molecules, causing C to T mismatches, as a measure of damage in sedaDNA^23^ using MetaDamage, as previously applied^9^. Interestingly, we found little difference in deamination signals between sediment types (ANOVA tests p=0.322, see Methods), or within sediment types over time, Fig. 2b. Cytosine deamination broadly correlates with age at a rate which is largely determined by regional environmental conditions of temperature and temperature fluctuation, but over shorter time periods stochasticity can be high^5^. If there is any contribution to DNA preservation through adsorption differences it is not of significant effect in this data set, satisfying the depositional model assumption that sediment type does not significantly influence DNA preservation and therefore contribution to taxonomic profile.

We further investigated the potential effects of habitat variation between samples, such as differences in salt concentration or pH, by categorizing samples on the basis of their taxonomic profiles previously determined^11^ into grassland, marsh, shallow marine or willow habitat (see Methods), Fig. 2c. As before, no significant difference occurs between habitat types in damage extent (ANOVA tests p=0.388, see Methods), suggesting that it is the regional environmental conditions of temperature and temperature fluctuation that are dominantly influential on DNA preservation^5^.

Over the whole dataset a weak trend occurs of increasing deamination signal over time (R^2^= 0.001, p=0.824, y=0.12 + 1.05E-6x ), but we detect no trends that would violate the underlying assumptions of the depositional model in this dataset.

Finally, we surveyed read frequencies of *Viridiplantae* assigned reads after stringent filtering which suggests that unstructured sand sediments (µ=8446, n=32) yield on average an order of magnitude more reads than unstructured silts (µ=418, n=21) in this data set, although the variance is generally wide between samples, Fig 2D. It may therefore be the case that an adsorbed sedaDNA fraction has remained adsorbed in the silts explaining the tendency to lower read counts recovered.

Laminated sediments of all types are associated with high read counts (sands µ=2234 n=36, silts µ=5619 n=101), suggesting that laminations, often comprised of organic material, represent rich sources of sedaDNA. Together, these data support the notion that sedaDNA profiles in this data set are largely based on free DNA rather than adsorbed DNA.

### Elucidation of an influx landscape based on sedimentation regime

The sediment influx depositional model consists of 8 parameters, *cl* (local contribution of sedaDNA)*, cs* (contribution of influxing sedaDNA from distal source)*, δ_xloc_* (amount of change observed due to local environmental change), *δ_xsed1_* (amount of change observed due to distal environmental change), *δ_xsed1-2_* (amount of change observed due to change in distal source), *id* (difference in distal environments associated with sedimentation change), *e_1_* (difference in extent of change of local relative to distal environments) and *e_2_* (change in contribution of influxing sources associated with change in sedimentation regime due to factors such as dynamic flow rates), Fig. 1. Together these parameters contribute to the difference observed in taxonomic profile when sediment regime remains unchanged (∂1) and when sedimentation regime changes (∂2) (see Methods). In order to understand how local and influxing sources of DNA are expected to change with varying values of ∂1 and ∂2 in this depositional model we estimated parameter landscapes for six variables (*cl*, *δ_xloc_*, *δ_xsed_*, *δ_xsed1-2_*, *e_1_* and *e_2_*), Fig. 3.

**Figure 3.**
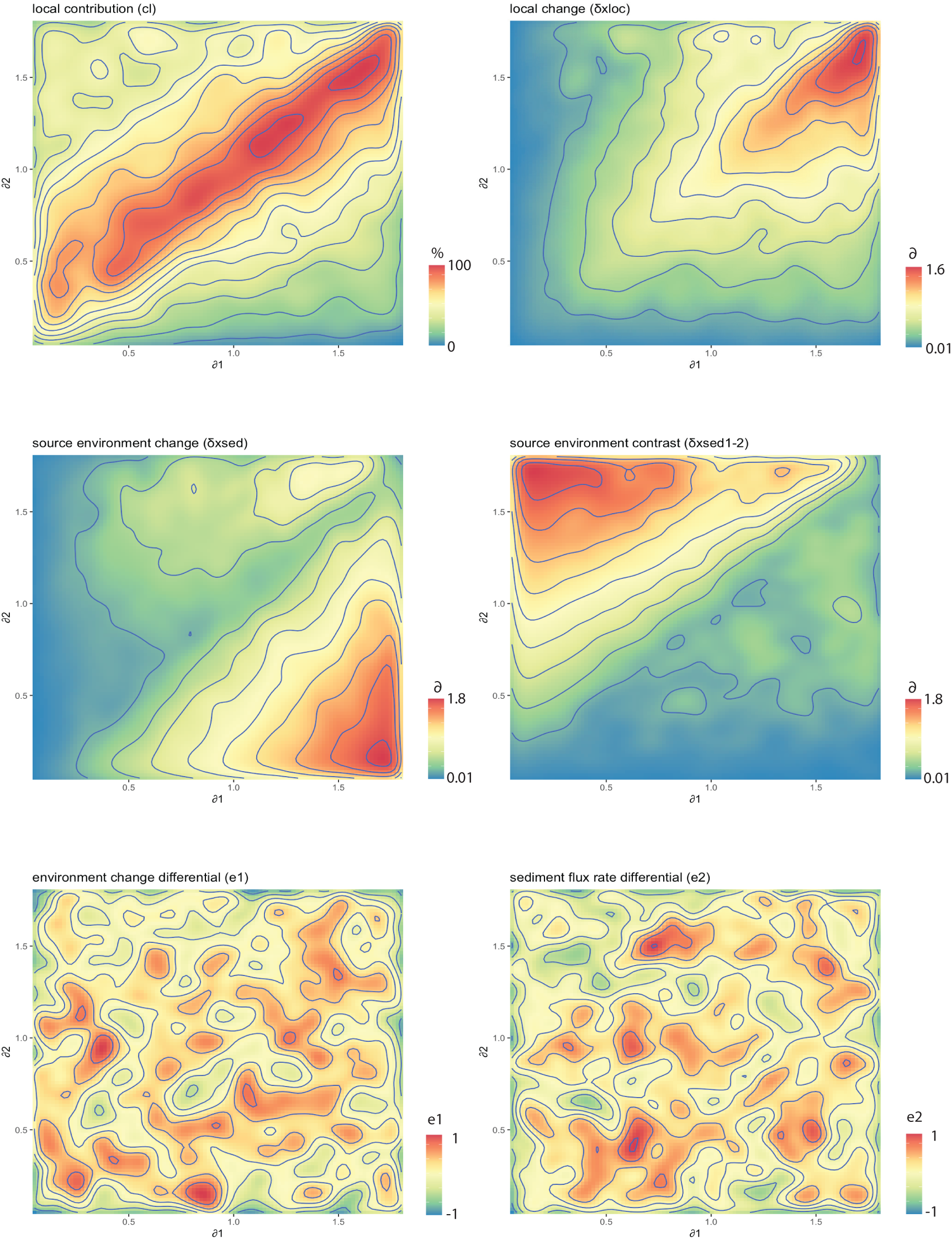
Parameter landscapes for sediment influx depositional model. Monte Carlo based solutions (see Methods) for model parameters *cl, δ_xloc_, δ_xsed_, δ_xsed1-2_, e1 and e2* for combinations of ∂1 and ∂2.

The landscape for local contribution (*cl*), shows that a high influence of local contribution is associated with situations in which the difference observed between taxonomic profiles is similar regardless of the sedimentation regime, Fig. 3a.

Intuitively, when both ∂1 and ∂2 are high, it is associated with a greater level of taxonomic change in the local environment, Fig 3b. However, a greater influence associated with influxing sediments is inferred when ∂1 and ∂2 become unbalanced, suggesting that changes to the sedimentation regime are influential on taxonomic profiles under these circumstances. When ∂1 is high relative to ∂2, it is associated with a greater level of taxonomic change in the distal environment from which the sediment is sourced. Conversely, when ∂2 is high relative to ∂1, it is associated with large changes in taxonomic profiles between changing sediment types, possibly suggesting likely changes in source location. These results suggest that even if sedaDNA may be inferred to have come from influxing sources and so not be appropriate for highly local interpretations, interpretations about the distal environment may be possible.

The two environmental variables *e_1_* and *e_2_* do not produce highly structured landscapes and so do not appear to be broadly informative. These variables account for the fact that the rate of biological change, in terms of taxa turnover, may vary between local and distal environments (*e_1_*), or that the contribution of influxing DNA may change with changing flow rates over time between distal source and local environments (*e_2_*). Neither of these variables lead to a systematic explanation of varying ∂1 and ∂2 values, even though they may themselves vary.

### Application of the taphonomy framework to different depositional systems

While the depositional model framework appears validated on the extensive alluvial data set from Doggerland^11^, it has yet to be applied to different ecological systems in which the taphonomic regime is expected to differ. The parameter landscapes shown in Fig.3 provide a look-up reference for any given values of ∂1 and ∂2. We leveraged this to examine published datasets of lacustrine^24–27^,permafrost^4,28^ and terrestrial^29,30^ systems in which there are sufficient sedimentological data to apply the depositional model. For each study, we calculated values of ∂1 and ∂2 for different sediment types (see Methods), and used these to plot coordinates in the *cl* ∂1/∂2 landscape (Fig. 3a) to give estimates of distal source contribution to sedaDNA profiles, Table 1. Despite the relatively low sample number on which ∂ values were based in many cases some informative trends emerged.

**Table 1.**
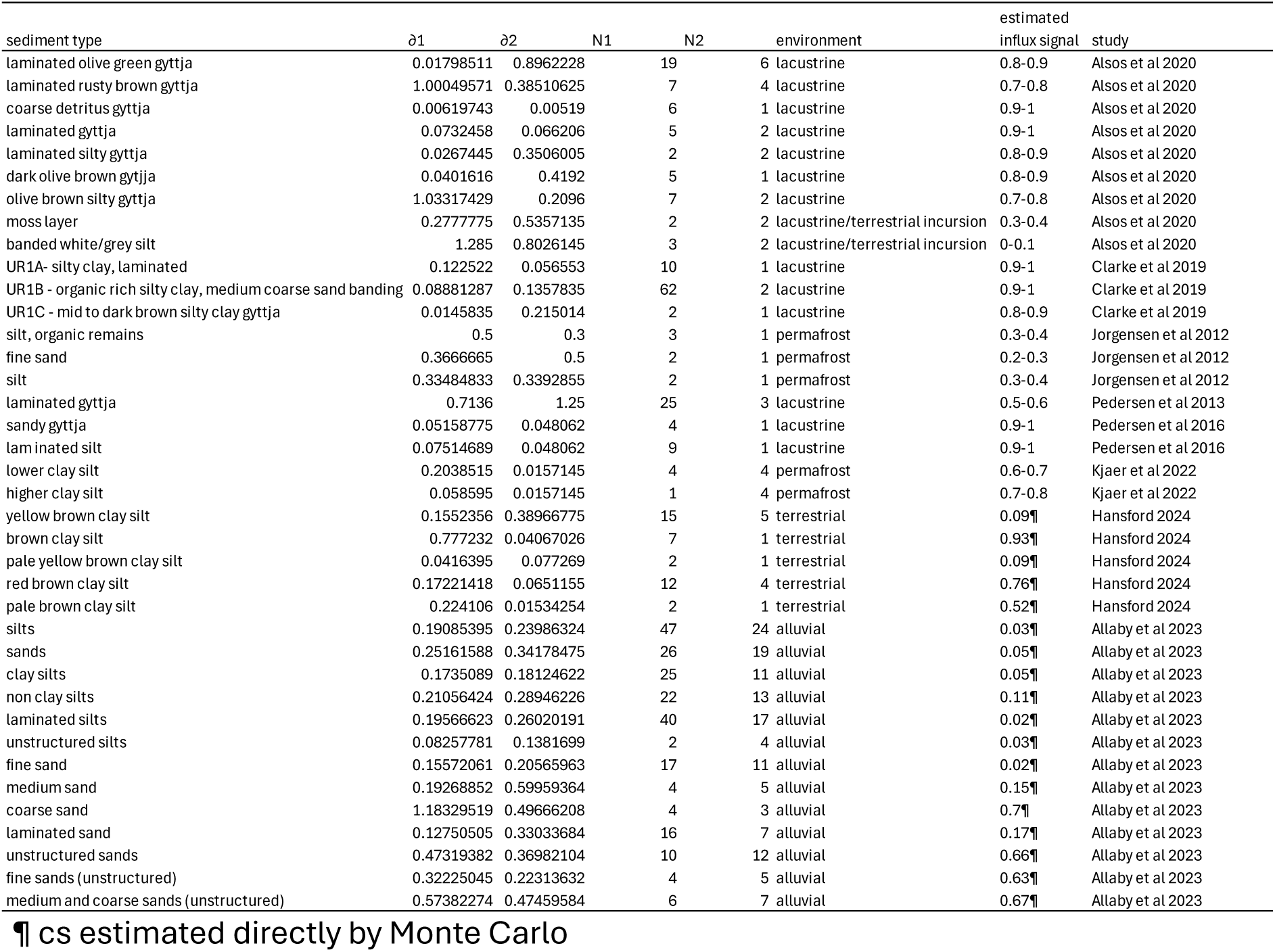
Influx signal (cs) estimates of published sedaDNA studies.

We examined 15 different sediment types of lake sediments across four studies^24–27^ (see Supporting Data 1-3). In all but three cases the ∂1/∂2 values were consistent with the majority of sedaDNA influxed from non-local regions in the depositional model, supporting previous findings of a regional sedaDNA signal from the catchment area for lake systems^12^. Two exceptions are sequential samples taken from a shallow area of the glacial lake Andøya in Norway (3 metres depth), dated to 14.18-12.73 kya^24^. The strata were exceptional in the profile because of the occurrence of a moss layer and underlying silts consistent with erosion of acidic mire from the lake edge^24^, which is reflected in the depositional model as a switch to local sources dominating the sedaDNA profile. The third example is from samples taken from Lake Comarum, a small (58 x 85 metres) shallow (> 1 metre) lake in southern Greenland^27^. The depositional model indicates an equable contribution to the sedaDNA profile from local and distal sources. Given the very small size of this lake it is not surprising that local vegetation in the immediate vicinity would have had a large influence on the sedaDNA profile leading to a mixed signal of local and distal sources.

We examined two permafrost studies. The first of these was based on three cores taken from around the Lake Tamyr area and three sediment types^28^ (see Supporting Data 4). In this case the depositional model solution indicates a domination of the sedaDNA signal from local sources, but an appreciable and consistent (25-35%) input from non-local sources, indicating the possibility of a mixed ecosystem signal. In this case, all cores were taken close to the lake edges, and while one might expect terrestrial systems to be dominated by a local signal in the absence of a strong external source of influx it is likely that periodic flooding could have introduced a more regional signal of sedaDNA over time. The second permafrost study was based on samples from Kap København formation in North Greenland^4^. (see Supporting Data 4). We found one suitable sequence of samples for analysis from sample site 69 (stratigraphic Unit B2), where a change in the silt sediments is apparent with a contrast in clay content indicating some change in sedimentation regime. In this case we estimate between 60 and 80% of the sedaDNA signal comes from distal sources leading to a likely mixed ecosystem signal, which matches the known taphonomy of the site of a terrestrial system washed into an estuary during an Early Pleistocene glacial minimum^31^.

While lacustrine and permafrost systems are usually assumed to have little post depositional DNA movement due to processes such as leaching^32^, terrestrial systems often show signs of post depositional DNA movement^8,33^. To mitigate this confounding issue, we examined terrestrial samples in which sedaDNA profiles had been tested statistically and no significant evidence of post depositional movement detected, which is further supported by a stable chemostrigraphy^29,30^. Samples were retrieved from the lower sections of Neolithic large man-made pits surrounding the settlement of Durrington Walls, below any instances of modern disturbance. In this case sediment profiles are all silts, which are graded by colour from yellow to red to brown. Yellow silts are consistent with a dominant local signal in the model. However, an increasing influence of distal sources is clearly correlated with progressively darker silt sediments, which for the most part overlay the lighter silts below. The current interpretation of chemostratigraphy and optically stimulated luminescence (OSL) profiles at this site is that the pits may have been infilled manually at later points in history after the Neolithic^30^. The depositional model suggests that this infilling was probably highly influential on the subsequent sedaDNA profile, and the sedaDNA itself appears to be highly associated with the source of colouration of the silts.

## Discussion

The depositional model is informative because of the dual nature of instances in which two critical variables are estimated, *cl* and *δ_xloc_*. While in the ∂1 scenario independent variables *cl*, *δ_xloc_* and *e_1_* are influential, in the alternative ∂2 it is independent variables *cl*, *δ_xloc_*, *id* and *e_2_* which are influential. Therefore, solutions of *cl* and *δ_xloc_* must satisfy both scenarios, consequently two independent sources of information are consulted analagous to simultaneous equations. The relative simplicity of the model facilitates its applicability to a wide range of environments without the overfitting effects of further peripheral parameters. The obvious further parameters that might be considered include the effects of various different sediment properties with regard to DNA adsorption, although we find no evidence that this has a confounding effect in our test dataset from Doggerland^11^. It may be the case that sedaDNA data sets are derived largely from unadsorbed, free DNA, although further investigation into the extent of deamination in different sediment types may refine understanding. Kjaer et al (4) observed differential recovery of DNA from model minerals with as little as 4% being recovered from the smectite associated fraction of sediments, but 43% is recoverable from quartz, suggesting differential mineralogical make up of sediments could result in the differential read count we observe between sandy and clay rich deposits. If DNA were released from a state of adsorption, the expected effect would be to influence the amount and level of damage of DNA present rather than the taxonomic profile as a whole. A more serious assumption of the model is the independence of sediment turnover and taxonomic turnover, that sediment changes do not necessarily reflect large events that also cause changes in taxonomic composition giving a spurious correlation through a common underlying variable. While it is unquestionably the case that some sediments reflect climatic conditions such as the loess of periglacial conditions^34,35^, on a local scale sediment turnover in our test data set is driven by more frequent ongoing local processes with no clear general correlation to larger climatic events such as the Younger Dryas or Pleistocene/Holocene boundary. This local dominance of turnover suggests that the geomorphological processes that the model tracks through sediment transitions are not expected to be directly linked taxonomic profile turnover. However, this may not be the case for all studies so should be routinely considered. In sum, the sedaDNA sediment influx deposition model leads to a taphonomy framework illustrated in Fig. 4.

**Figure 4.**
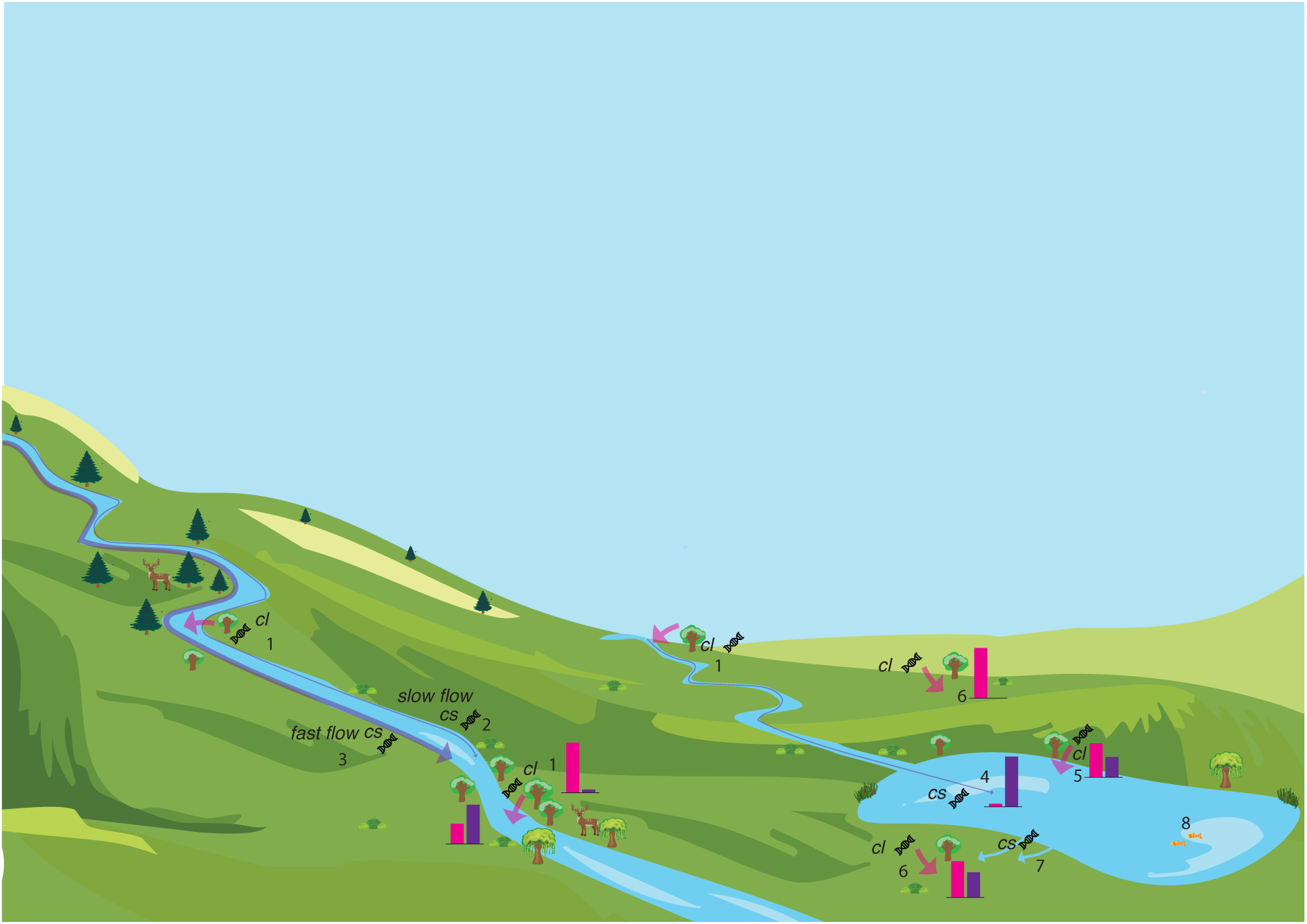
DNA deposition patterns based on sediment influx depositional model outputs from observed sedaDNA data. Arrows show direction of DNA movement, arrow thickness represents quantity of DNA. Bar charts indicate relative contribution of local (pink) and distal (purple) sedaDNA sources. 1. Local contribution (cl) substantial in alluvial systems. 2. Little DNA is influxed (cs) in slow moving water due to DNA removal processes. 3, Substantial DNA can be influxed in rapidly moving water. 4. Lacustrine systems mostly receive influxed DNA with few local sources of DNA resulting in a dominant influx signal, even if little DNA is involved. 5. Lake margins can supply substantial local source DNA in the immediate vicinity. 6. Terrestrial systems are expected to be dominated by local sources in the absence of an influx source. 7. Events such as flooding can provide an influx source of DNA. 8. Faunal sources in lacustrine systems could potentially provide a dominating local source of DNA.

### Alluvial and Lacustrine systems have contrasting sedaDNA taphonomies

The depositional model originally identified an association in alluvial systems of slow moving (silt and clay associated) systems with a dominance of locally sourced sedaDNA signal with little or no sedaDNA arriving with sediments^11^. However, lake profiles of equably fine sediments show that sedaDNA originates from distal sources, Table 1. The half-life of DNA in fresh water is less than an hour^6^, which should seemingly apply in both taphonomic contexts, alluvial and lacustrine. From this we conclude that the amount of sedaDNA arriving from distal sources is likely to be comparable between the two and the difference between the estimates of local versus distal contributions must be in the absolute amount of DNA coming from local sources. A corollary is that lacustrine sedaDNA profiles are typically expected to be based on a lower absolute amount of DNA than alluvial profiles. It is intuitive that lakes provide little direct plant DNA from local sources if they are too deep for aquatic plants. However, it may be the case that a different taphonomic profile may be obtained from animal DNA than plant DNA in lacustrine systems where substantial local contribution may be made by fish and aquatic invertebrates.

### The prevalence of mixed ecosystem signals

Where distal sources of DNA are dominantly influential on the sedaDNA profile such as in lacustrine sediments the signal is likely to reflect a regional catchment signal, which is likely to include a range of ecosystem types. Where there is a large input of DNA from the immediate vicinity relative to any influxing material, as is expected to be the case for terrestrial systems, slow moving narrow river systems and very small bodies of water, the signal is likely to strongly reflect the immediate ecosystem.

However, there are numerous instances shown here in which there is evidence of input from both local and distal sources, with neither completely dominating. In such instances caution needs to be applied when making inferences about the local environment, or indeed the regional profile since the signal will be distorted by the type of local environment. It is therefore likely that many current published data sets represent mixed ecosystem signals, and some reappraisal may be appropriate. This is particularly the case for systems in which there has been past fast influxing flow, typically leading to medium to coarse sandy unstructured deposits. Where a large number of sedimentological shifts can be sampled in studies more robust estimates of deposition sources will be possible.

### Sediment reworking signals can be revealed under some circumstances

Where sedaDNA profiles can be attributed to distal sources it is possible that the DNA can reflect the contemporaneous distal environment or potentially reworked sedaDNA from an earlier time. If such reworked deposits are not vastly different in age, for example by only a few thousand years, it is unlikely that reworking would be discernible on the basis of clearly greater damage signatures in the older DNA given the stochasticity in damage, as apparent in Fig. 2b. However, sedaDNA that is of significant age, such as over half a million years, is typically associated with high frequencies of observed C to T mismatches indicating saturation of signal^4,36,37^ and so could be discernible from the unsaturated Holocene to late Pleistocene aged associated damage as outlined in Fig 2b. In such cases it may be possible to infer a reworking signal. Alternatively, reworking may be inferred where a sedaDNA profile attributed to an influxing signal closely resembles an earlier likely source. This was the case in the Southern River system in which sandy deposits identified as an influxing signal in core 31 showed marshy grassland profile in contrast to underlying locally sourced deposits. However, upstream almost identical taxonomic profiles of local source were identified a few hundred years earlier in age providing convincing evidence of a reworking episode^11^.

### Concluding remarks

The sediment influx depositional model is a potentially powerful tool to aid interpretation of sedaDNA profiles and highlights the need for comprehensive sedimentological data to be included in sedaDNA studies. In particular, the pervasive nature of mixed ecosystem signals combined with the current limits in identifying reworked signals underscores the caution required in interpretations and the need to be able to identify where profiles represent local or distal profiles.

## Methods

### Environmental correlation analysis

The environmental correlation analysis was carried using data from reference (11). All analyses were carried out using sedaDNA samples with filtered read counts that exceeded 300. The rate of environmental change between adjacent samples in a core (Fig 2a) was calculated using the difference in ecological distances (∂ values) between two consecutive samples (table S8 ^11^), divided by the difference in age of the samples. To calculate the rate of sediment change (Fig 2a), samples were ordered chronologically within cores and instances where sediment remained constant were given a value of 0 and given a value of 1 where the sediment changed. The number allocated to the sediment change was then divided by the difference in ages between the two samples to obtain a value for the rate of sediment change. A sliding window average approach was used for both sediment and biotic rates of change where the rate was dated to the mid-point range of the two sample ages, values were placed in date order and a sliding average of 4 samples was used to calculate the rate over time. The amount of damage present for different samples was determined using observed C to T mismatch frequencies at position 0 from MetaDamage^9^.

Samples were categorized by sediment type (Fig 2B) and habitat (Fig 2C) to allow comparisons of observed C to T mismatch frequencies at position 0 between different sediment and habitat types over time. Habitat types were determined using the dominant (>70%) ecosystem profile type from reference (11), whilst sediment types were determined using Fig S2^11^. Violin plots were generated in R^38^ using total *Viridiplantae* read counts of samples, separated into their sediment types (Table S2^11^).

### Basic model outline

The model outlined in Fig. 1 is based on differences in taxonomic profile (∂) that are measured as a difference in the sum of squares (*dss*) statistic of proportions of sedaDNA read profiles belonging to each taxonomic unit:

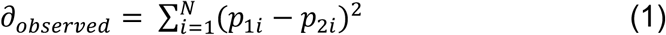

Where there are *N* taxonomic units, *p_1i_* and *p_2i_* are the proportions of the *i*th taxonomic unit in samples 1 and 2 respectively. Observed values of ∂1 and ∂2 were estimated from the average of all observed instances of adjacent samples within the same unchanged sediment type (∂1), and adjacent samples spanning sediment transitions (∂2). There is a general assumption that the mean time difference between samples in these two scenarios does not significantly differ reflecting a relatively even sampling regime.

The model difference expected under ∂1 is expressed as the sum of the amount of change due local environmental change (*δ*_xloc_ ) and amount of change due to distal environmental change associated with sedaDNA influxing with sediment (*δ*_xsed1_):

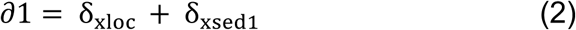

Where the amount of change attributable to influxing sedaDNA is a function of the relative proportional contributions of *δ*_xloc_ (*cl*) and *δ*_xsed1_ (*cs* where *cs =* 1-*cl*) and the rate of environmental change in the distal environment relative to the local environment (*e_1_*):

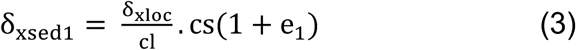

The model difference expected under ∂2 is expressed as the sum of the amount of change due local environmental change (*δ*_*xloc*_) and amount of change due to and the change associated with the compositional difference between sediments 1 and 2 (*δ_xsed1-2_*):

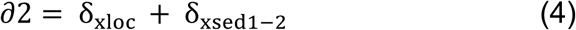

Where the amount of change attributable to influxing sedaDNA (*δ*_*xsed1-2*_) is a function of the change of the taxonomic profiles between sediments 1 and 2 (id), the relative proportional contribution of incoming sedaDNA (*cs*) and the associated change in that contribution associated with changing flow rates associated with the different sediment types (*e_2_*):

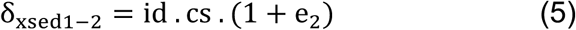

### Generation of parameter landscapes

In order to estimate parameter values from the model from observed data we used a Monte Carlo approach. For each pair of values of *e_1_* and *e_2_* there are three independent variables *cl*, *δ*_*xloc*_ and *id,* from which all other values are derived. Monte Carlo chains began with random values of *e_1_* and *e_2_* in the range of −1 to +1, followed by random increments of 0.01, for which all values of *cl*, *δ*_*xloc*_ and *id* were explored in increments of 0.01 in the ranges of 0-1, 0-2 and 0-2 respectively. For each set of parameter values model values of ∂1 and ∂2 were calculated using equations 2–4, and the score of the fit to observed values of ∂1 and ∂2 as follows:

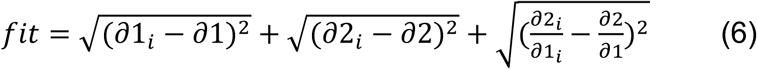

Unheated chains were terminated after 20 attempts to improve fit had failed and chains were repeated 1000 times. The entire parameter space is 1.6 x 10^11^, and chains were sufficient to explore most if not all this space.

In order to generate the landscapes parameters were estimated as described above for values of ∂1 and ∂2 ranging from 0.05 - 1.8 in 0.05 increments. Density plots with contours were generated using ggplot in R^39^.

### Estimation of ∂1 and ∂2 values from published data sets

In the case of lacustrine^24–27^ and permafrost^4,28^ data sets, the taxonomic unit used to calculate ∂ values was the species. The unit in the case of the terrestrial^29,30^ and alluvial^11^ systems was the guild unit, a group of species shown to co-occur in the data (see reference 11 for details). Authors’ measurements (read count, or metabarcoding presence) were converted into proportions of all observations for each sample, from which dss statistics were calculated. All workings are shown in Supplementary Data 1-5.

## Acknowledgements

This was supported by the NERC CENTA PhD program (NE/S007350/1), European Research Council (ERC) under the European Union’s Horizon 2020 research and innovation programme (ERC funded project No. 670,518 ‘Europe’s Lost Frontiers Project’, https://europa.eu/european-union/index_en, https://lostfrontiers.teamapp.com/), with continuing support from the UKRI Natural Environment Research Council (NERC) (NE/W005751/1) and the UKRI Arts and Humanities Research Council (AHRC) (AH/W003287/1).

## References

1. Orlando L, Allaby RG, Skoglund P, Der Sarkissian C, Stockhammer PW, Avila-Arcos M, Fu Q, Krause J, Willerslev E, Stone AC, Warinner C (2021) Ancient DNA Analysis. Nature Reviews Methods Primers 1:14

2. Rawlence, N. J. et al. Using palaeoenvironmental DNA to reconstruct past environments: Progress and prospects. J Quat Sci 29, 610–626 (2014).

3. Willerslev, E. et al. Ancient Biomolecules from Deep Ice Cores Reveal a Forested Southern Greenland Science 317, 111–114 (2007).

4. Kjaer, K. et al. A 2-million-year-old ecosystem in Greenland uncovered by environmental DNA. Nature 612, 283–291 (2022).

5. Kistler, L., Ware, R., Smith, O., Collins, M., Allaby, R. A new model for ancient DNA decay based on paleogenomic meta-analysis. Nucleic Acids Res. 45, 6310–6320 (2017).

6. Seymour M et al Acidity promotes degradation of multi-species environmental DNA in lotic mesocosms. Comm. Bio. 1, 4 10.1038/s42003-017-0005-3 (2021).

7. Sand, K. K., Jelavić, S., Kjær, K. H. & Prohaska, A. Importance of eDNA taphonomy and provenance for robust ecological inference: insights from interfacial geochemistry. Environmental DNA 6:3519

8. Haile, J., et al. Ancient DNA Chronology within Sediment Deposits: Are Paleobiological Reconstructions Possible and Is DNA Leaching a Factor? Mol. Biol. Evol. 24, 982–989 (2007).

9. Gaeney, V., et al. Multi-proxy characterisation of the Storegga tsunami and its impact on the early Holocene landscape of the southern North Sea. Geosciences 10, 270 (2020).

10. Ambrecht, L.H., et al. Ancient DNA from marine sediments: Precautions and considerations for seafloor coring, sample handling and data generation. Earth-Science Reviews 196, 102887 (2019).

11. Allaby, R. et al. Pleistocene-Holocene sedaDNA reconstruction of Southern Doggerland reveals early colonization before inundation consistent with northern refugia. Research Square (2023) doi:10.21203/RS.3.RS-3296992/V1

12. Giguet-Covex, C. et al. New insights on lake sediment DNA from the catchment: importance of taphonomic and analytical issues on the record quality. Scientific Rep. 9,14676 (2019).

13. Edwards, M. E., et al. Metabarcoding of modern soil DNA gives a highly local vegetation signal in Svalbard tundra. Holocene 28, 2006–2016 (2018).

14. Jorgenson T et al. A comparative study of ancient sedimentary DNA, pollen and macrofossils from permafrost sediments of northern Siberia reveals long-term vegetational stability. Mol. Ecol. 21, 1989–2003 (2012).

15. Pedersen MW et al. A comparative study of ancient environmental DNA to pollen and macrofossils from lake sediments reveals taxonomic overlap and additional plant taxa. Quat. Sci. Rev. 75,161e–168 (2013)

16. Parducci, L., Alsos, I.G., Unneberg, P., Pedersen, M.W., Han, L., Lammers, Y., Salonen, J.S., Väliranta, M.M., Slotte, T., and Wohlfarth, B. Shotgun Environmental DNA, Pollen, and Macrofossil Analysis of Lateglacial Lake Sediments From Southern Sweden. Frontiers Ecol. Evol. 7, 189 (2019).

17. Alley, R.B., Ágústsdóttir, A.M. The 8k event: cause and consequences of a major Holocene abrupt climate change. Quaternary Science Reviews 24,1123–1149 (2005).

18. Pedreira-Segade, U. et al. How do nucleotides adsorb onto clays? Life 8, 7–8 (2018).

19. Yu, W. H. et al. Adsorption of proteins and nucleic acids on clay minerals and their interactions: A review. Appl Clay Sci 80–81, 443–452 (2013).

20. Cai, P., Huang, Q., Zhang, X. & Chen, H. Adsorption of DNA on clay minerals and various colloidal particles from an Alfisol. Soil Biol Biochem 38, 471–476 (2006).

21. Cai, P., Huang, Q., Zhang, X. Microcalorimetric studies of the eeects of MgCl2 concentrations and pH on the adsorption of DNA on montmorillonite, kaolinite and goethite. Appl Clay Sci 32, 147–152 (2006).

22. Jelavic, S., Lanson, B., Findling, N., Lanson, M., A route for long-term DNA preservation through nanoconfinement in smectites. ChemRxiv., doi:10.26434/chemrxiv-2023-6m43n (2023).

23. Brotherton, P., Endicott, P., Sanchez, J.J., Beaumont, M., Barnett, R., Austin, J., Cooper, A. Novel high-resolution characterization of ancient DNA reveals C’U-type base modification events as the sole cause of post mortem miscoding lesions. Nucleic Acids Res. 35, 5717–5728 (2007)

24. Alsos, I.G. et al. Last Glacial Maximum environmental conditions at Andøya, northern Norway; evidence for a northern ice-edge ecological “hotspot”. Quaternary Sci. Rev. 239, 106364 (2020).

25. Clarke, C.L., et al. Holocene floristic diversity and richness in northeast Norway revealed by sedimentary ancient DNA (sedaDNA) and pollen. Boreas 48, 299–316 (2019).

26. Pederson, M.W., et al. Postglacial viability and colonization in North America’s ice-free corridor. Nature 537, 45–49 (2016).

27. Pedersen, M.W., et al. A comparative study of ancient environmental DNA to pollen and macrofossils from lake sediments reveals taxonomic overlap and additional plant taxa. Quaternary Sci. Rev. 75, 161–168 (2013).

28. Jørgensen, T., et al. A comparative study of ancient sedimentary DNA, pollen and macrofossils from permafrost sediments of northern Siberia reveals long-term vegetational stability. Mol. Ecol. 21, 1989–2003 (2012).

29. Hansford, T. SedaDNA analysis of Durrington Walls: a new class of Neolithic monument. PhD thesis, University of Warwick (2024).

30. Gaeney, V. et al. The Perils of Pits: further research at Durrington Walls henge (2021– 2023). Internet Archaeology XXXXX (2025).

31. Funder, S. et al. Late Pliocene Greenland—the Kap København formation in North Greenland. Bull. Geol. Soc. Den. 48, 117–134 (2001).

32. Parducci, L. et al. Ancient plant DNA in lake sediments. New Phytologist 214, 924– 942 (2017).

33. Andersen, K., et al. Meta-barcoding of ‘dirt’ DNA from soil reflects vertebrate biodiversity. Mol. Ecol. 21, 1966–1979 (2012).

34. Pye, K. The nature, origin and accumulation of loess. Quaternary Sci. Rev. 14, 653–667.

35. Frechen, M. Loess in Europe. Quaternary Sci. J. 60, 3–5.

36. Massilani, D., et al. Microstratigraphic preservation of ancient faunal and hominin DNA in Pleistocene cave sediments. Proc. Natl. Acad. Sci. U.S.A. 119, e2113666118 (2022).

37. Slon, V. et al. Neandertal and Denisovan DNA from Pleistocene sediments. Science 356, 605–608 (2017).

38. 38. R Core Team (2021). R: A language and environment for statistical computing. R Foundation for Statistical Computing, Vienna, Austria. URL https://www.R-project.org/.

39. Wickham H. ggplot2: Elegant Graphics for Data Analysis. Springer-Verlag New York (2016).

